# Generation of thymidine auxotrophic *Agrobacterium tumefaciens* strains for plant transformation

**DOI:** 10.1101/2020.08.21.261941

**Authors:** Ephraim Aliu, Mercy K. Azanu, Kan Wang, Keunsub Lee

## Abstract

*Agrobacterium*-mediated transformation is a widely used gene delivery method for fundamental researches and crop trait improvement projects. Auxotrophic *Agrobacterium tumefaciens* strains are highly desirable for plant transformation because they can be easily removed from the explants after co-cultivation due to their dependence on essential nutrient supplementation. The thymidine auxotrophic *A. tumefaciens* strain LBA4404Thy- has been successfully used for plant transformation, however, auxotrophic version of other commonly used strains are not available yet to public laboratories. Here we report the generation of EHA101, EHA105 and EHA105D thymidine auxotrophic strains. These strains exhibited thymidine-dependent growth in the bacterial medium, and the transient GUS expression assay using Arabidopsis seedling showed that they retain the equivalent T-DNA transfer capability as the original strains thus are suitable for plant transformation.

## Introduction

*Agrobacterium tumefaciens* is a widely used plant biotechnology tool for plant genome editing and crop improvement (Gelvin 2003). The unique ability of *A. tumefaciens* to transfer its own DNA (T-DNA) into plant genome has been widely utilized to efficiently deliver a low copy number of transgenes into plant genomes (Gelvin 2003; Tzfira and Citovsky, 2006). Virulence (*vir*) genes encoded on the tumor inducing (Ti) plasmid and chromosome (*chv* genes) enable *A. tumefaciens* strains sense the environmental signals exuded from the wounded plants, such as low pH, simple sugars and phenolic compounds to initiate T-DNA delivery process (Gelvin 2003; Gelvin 2017). Natural T-DNAs carry the biosynthesis genes for plant hormones to induce the crown gall tumor formation and the opine biosynthesis genes to produce the small molecular weight compounds that are used as important energy source by the infecting *A. tumefaciens* (Tempé & Petit 1982; Britton et al., 2008). T-DNAs are flanked by the two short repeat sequences, i.e., left and right borders (LB and RB), and a gene of interest can be provided *in trans* on a binary vector (Lee and Gelvin, 2008).

For plant transformation purposes, *A. tumefaciens* strains have been engineered for several aspects: first, the Ti plasmids were ‘disarmed’ by deleting the natural T-DNAs (Ooms et al., 1981; Hood et al., 1986; Koncz et al., 1986). Second, the supervirulent Ti plasmid pTiBo542 (Komari et al., 1986) was transferred to different chromosomal backgrounds including C58 resulting in highly virulent *Agrobacterium* strains such as A281 (Sciaky et al., 1978). Third, extra *vir* genes were provided on a ‘super-binary vector’ (Komari et al., 2006) or a ‘helper plasmid’ (Anand et al., 2018) to further enhance transformation frequency. Additionally, *recA* recombinase gene was inactivated to reduce unwanted plasmid DNA rearrangements within the *Agrobacterium* cells (Lazo et al., 1991) and more recently, a thymidine auxotrophic strain (LBA4404Thy-) was generated and successfully used for plant transformation (Ranch et al., 2010; Anand et al. 2018).

The thymidine auxotrophic LBA4404Thy-strain was generated by deleting the thymidylate synthase gene (*thyA*) and it cannot survive without supplementing thymidine in the medium (Ranch et al., 2010). Thymidine auxotrophic *Agrobacterium* cells can be easily removed from the explants after the co-cultivation period without using antibiotics; thus it is more economical and helpful to avoid antibiotic toxicity to delicate plant tissues (Pollock et al., 1983; Nauerby et al., 1997). Moreover, the *Agrobacterium* strains that carry the reagents for plant genome engineering, such as clustered regularly interspaced palindromic repeats (CRISPR) systems and herbicide resistance genes, are much less likely to survive in the natural environments, easing some of the biosafety concerns. Therefore, auxotrophic *Agrobacterium* strains are highly desirable for plant transformation; however, the auxotrophic version of the most commonly used public *Agrobacterium* strains, such as EHA101 and EHA105, are not available yet. Here, we report the generation of thymidine auxotrophic *Agrobacterium* strains of EHA101, EHA105 and an EHA105 derivative EHA105D (Lee et al., 2013) by homologous recombination-mediated *thyA* gene knockout. These auxotrophic strains were not able to grow in the bacterial medium without thymidine supplementation and more importantly, they retained the T-DNA transfer capabilities hence can be used for plant transformation.

## Materials and Methods

### Bacterial strains and growth conditions

*Agrobacterium tumefaciens* strains, plasmids, and primer sequences used in this study are listed in Tables 1 and 2. LBA4404 thymidine auxotrophic strain (LBA4404Thy-) was obtained from Corteva Agriscience (Ranch et al., 2010). Three *Agrobacterium tumefaciens* strains EHA101 (Hood et al., 1986), EHA105 (Hood et al., 1993) and EHA105D (Lee et al., 2013) were used to generate thymidine auxotrophs via homologous recombination-mediated *thyA* knockout. These three strains have the *A. tumefaciens C58* chromosomal background. EHA105D was derived from EHA105 by deleting *atsD* gene (*Atu5157*), which might play a role for *Agrobacterium* attachment to plant cells (Matthysse et al., 2000). EHA105D showed slightly higher maize transformation frequency than EHA105 using the binary vector pTF101.1 (Lee et al., 2013).

**Table 1:** Strains and plasmids used in this study.

| Strains and plasmids | Description | Reference |
| --- | --- | --- |
| **Strains** |  |  |
| LBA4404 Thy- | *thyA* knockout mutant used as a positive control for AGROBEST assay | Ranch et al. 2010 |
| EHA101 | derivative of A281 (A136/pTiBo542), kanamycin resistant | Hood et al. 1986 |
| EHA101 Thy- | *thyA* knockout mutant derived from EHA101 | This study |
| EHA105 | derivative of A281 (A136/pTiBo542) | Hood et al. 1993 |
| EHA105 Thy- | *thyA* knockout mutant derived from EHA105 | This study |
| EHA105D | *atsD* knockout mutant derived from EHA105 | Lee et al. 2013 |
| EHA105D Thy- | *thyA* knockout mutant derived from EHA105D | This study |
| **Plasmids** |  |  |
| pTFsacB | Suicide vector derived from pK19mobsacB (Schäfer et al. 1994) | This study |
| pKLSacB | Suicide vector derived from pTFsacB with spectinomycin resistance gene | This study |
| pMKA1 | *thyA* knockout vector with *sacB* and kanamycin resistance gene | This study |
| pKL2128 | *thyA* knockout vector with *sacB* and spectinomycin resistance gene | This study |
| pTF102 | Binary vector with a *gus* gene | Frame et al. 2002 |

**Table 2:**
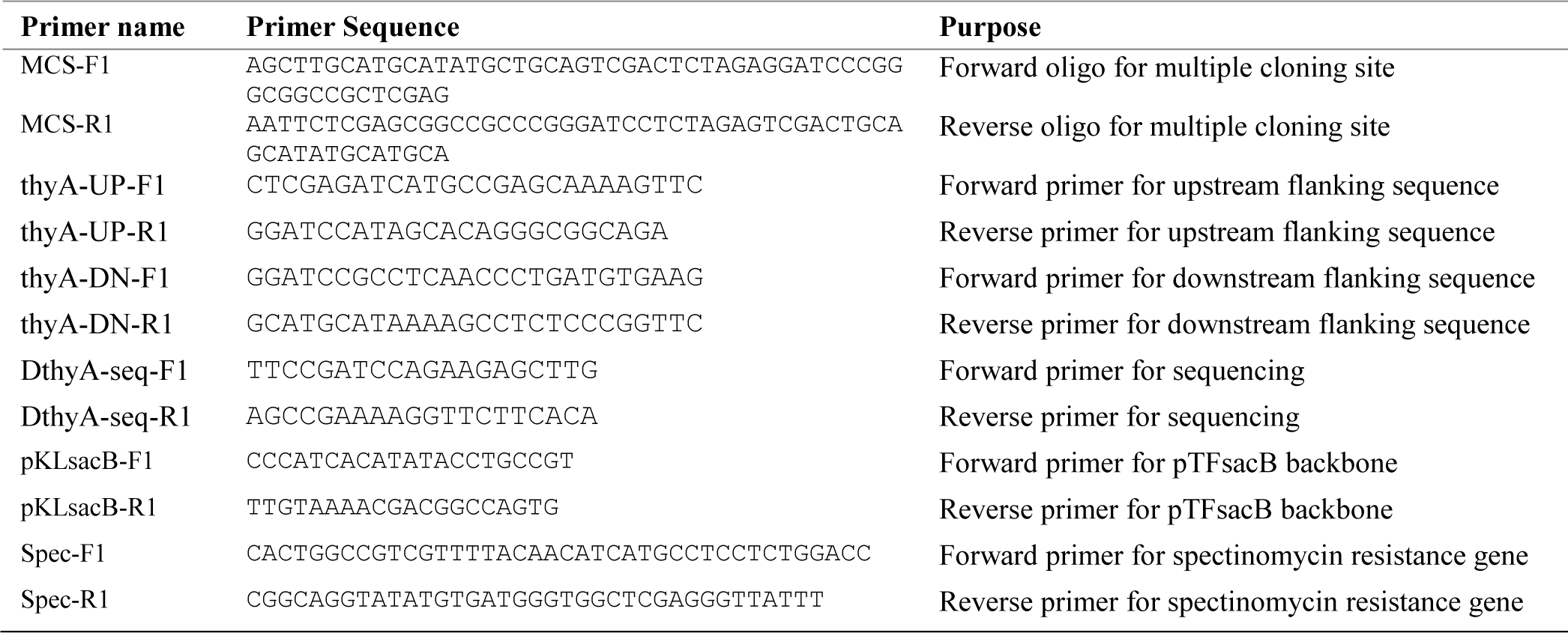
Primers used in this study.

Agarose gel electrophoresis, restriction enzyme digestion, and other molecular techniques employed were conducted according to standard protocols or as stated in the manufacturer’s instructions. Small amounts of plasmid DNA were prepared using QIAprep spin Miniprep Plasmid Kit (Qiagen, Hilden, Germany). DNA purification from agarose gel was done using the QIAEX II gel extraction kit (Qiagen). Restriction enzymes were purchased from the New England Biolabs (MA, USA) and oligonucleotides were synthesized by Integrated DNA Technologies (Iowa, USA). Standard Sanger sequencing analyses were performed by the DNA Facility at the Iowa State University (Iowa, USA).

### Vector construction

Knockout constructs were made as previously described (Ranch et al., 2010; Lee et al., 2013) using a *sacB*-based suicide vector (Schäfer et al., 1994). Firstly, pTFsacB was made by replacing the multiple cloning site (MCS) of pK19mobsacB (Schäfer et al., 1994). pK19mobsacB was digested with HindIII and EcoRI and then ligated with the annealed oligonucleotides MCS-F1 and MCS-R1 (Table 2). Secondly, PCR primers were designed using the Primer3 software (Untergasser et al., 2012) to amplify about 800 bp of upstream (thyA-UP) and downstream (thyA-DN) flanking sequences of *thyA* (*Atu2047*) from *A. tumefaciens* strain C58. PCR amplification was performed using the Phusion high-fidelity DNA polymerase (ThermoFisher Scientific, MA, USA) and *A. tumefaciens* C58 genomic DNA according to the manufacturer’s instruction. A 20 µl PCR reaction mix included 1X Phusion HF buffer, 125 µM dNTPs, 0.5 µM primers, and 0.4 unit of Phusion high fidelity DNA polymerase. Thermocycling conditions were as follows: initial denaturation for 30 sec at 98 °C, followed by 30 cycles of 10 sec at 98 °C, 15 sec at 63 °C, 30 sec at 72 °C, and a final extension for 5 min at 72 °C. thyA-UP and thyA-DN PCR products were then individually cloned into pJET1.2 cloning vector (ThermoFisher Scientific, MA, USA). Sanger sequencing analyses verified the flanking sequences before they were digested with XhoI/BamHI and BamHI/SphI, respectively. pTFsacB was digested with XhoI/SphI and ligated with thyA-UP and thyA-DN fragments to produce the knockout construct, pMKA1 (Figure S1A). Because EHA101 has kanamycin resistance (Hood et al., 1986), pMKA1 could not be used. Thus, we replaced the kanamycin resistance gene in pTFsacB with a spectinomycin resistance gene to make pKLsacB. pTFsacB backbone excluding the kanamycin resistance gene cassette was PCR amplified using pKLsacB-F1 and R1 (Table 2) and assembled with the spectinomycin resistance gene cassette amplified with Spec-F1 and R1 using pYPQ141D (Lowder et al., 2015) as a template. NEBuilder Hifi DNA assembly cloning kit (New England BIolabs, MA, USA) was used for the final assembly. pKLsacB was then digested with XhoI/SphI and ligated with the thyA-UP/thyA-DN fragment cut from pMKA1, resulting in pKL2128 (Figure S1B). pMKA1 was used for EHA105 and EHA105D, while pKL2128 was used for EHA101 to generate thymidine auxotrophic strains.

### Generation of *thyA* knockout mutants

pMKA1 and pKL2128 (Figure S1) were introduced into *Agrobacterium* strains EHA101, EHA105 and EHA105D by electroporation as described previously (Mattanovich et al., 1989) using a Bio-Rad Gene Pulser (Bio-Rad, CA, USA). After electroporation, 500 µL of SOC medium was added and incubated in a 28°C incubator for 2 hrs with shaking at 200 rpm. *Agrobacterium* cells were collected by centrifugation and resuspended in about 100 µL of SOC medium and spread on fresh Yeast Extract Peptone (YEP, Table S1) plate amended with appropriate antibiotics. Plates were sealed with parafilm and incubated at 28°C for two days. Antibiotic-resistant colonies (kanamycin-resistant EHA105 and EHA105D; spectinomycin-resistant EHA101) were picked and resuspended in 500 µL of fresh YEP medium in 1.5 mL microcentrifuge tubes and 100 µL was spread on solid YEP medium amended with 5% sucrose and 150 mg/L of thymidine. Two days later, well-isolated colonies were picked and inoculated on three plates: YEP without thymidine, YEP with 50 mg/L thymidine, and YEP with 50 mg/L thymidine and kanamycin (50 mg/L) or spectinomycin (100 mg/L). Colonies that can grow only on YEP with 50 mg/L thymidine plate were PCR screened using primers DthyA-seq-F1 and DthyA-seq-R1 (Table 2), and about 274 bp PCR products were subjected to Sanger sequencing to verify precise *thyA* deletion mutants.

### Evaluation of T-DNA delivery ability of thymidine auxotrophic strains

We used AGROBEST assay (Wu et al., 2014) to test if the *thyA* deletion mutants retain T-DNA delivery capability. *Arabidopsis thaliana* T-DNA insertion mutant *efr-1* (SALK 044334) was obtained from the Arabidopsis Biological Resource Center (Columbus, Ohio). About 300-500 seeds were surface sterilized in a 1.5 mL tube by soaking in 1 mL of 50% bleach (3% sodium hypochlorite) and 0.1% SDS solution for 15 mins followed by rinsing four times with sterile water. One milliliter of 1/2 MS medium supplemented with 5% sucrose was added to each tube, and seeds were transferred to a 60 mm petri dish using a wide-bore pipette tip and a pipette. The *efr*-1 seeds were subjected to a cold treatment (4°C) for 48 hrs for synchronized seed germination and grown for 7 days in a growth chamber at 22°C under a 16 hr/8 hr light/dark cycle.

Thymidine auxotrophic strains and their corresponding wildtype strains (positive control) were transformed with the binary vector pTF102 (Frame et al., 2002) by electroporation as described above. The binary vector pTF102 carries a GUS reporter gene (ß-glucuronidase) driven by a cauliflower mosaic virus 35S promoter. *Agrobacterium* strains were grown for 20 hrs in 5 mL of YEP medium supplemented with 50 mg/L thymidine (for auxotrophs) and appropriate antibiotics (50 mg/L kanamycin and 100 mg/L spectinomycin for EHA101; 100 mg/L spectinomycin for EHA105 and EHA105D) in 50 mL tubes at 28°C with 200 rpm. Immediately before infection, *Agrobacterium* cells were pelleted by centrifugation and re-suspended in AB induction medium (Gelvin 2006) to a density of OD_550_ = 0.04.

For *Agrobacterium* infection, about 10 *Arabidopsis* seedlings were transferred to each well of a 12-well plate using sterile inoculation loops. To each well, 500 µL of 1/2 MS medium (Table S1) was aliquoted before adding equal volume (500 µL) of freshly prepared *Agrobacterium* cell suspensions supplemented with 50 mg/L thymidine. Each *Agrobacterium* strain was added to three wells (replicates) in each experiment. The 12-well plates were sealed with 3 M micropore tape and incubated in a growth chamber for two days at 22°C under a 16 hr/8 hr light/dark cycle. After two-day co-cultivation, *Agrobacterium* cells were removed by pipetting and one milliliter of fresh 1/2 MS medium amended with 100 mg/L cefotaxime and 100 mg/L timentin were added into each well and *Arabidopsis* seedlings were further grown for two days in the growth chamber. EHA105 strain without pTF102 was used as negative control.

Transient transgene expression was visualized by GUS staining as previously described with slight modifications (Cervera 2005). Briefly, liquid medium was removed, and 1 mL of GUS staining solution was added to Arabidopsis seedlings in a 12-well plate and incubated at 37°C overnight. Following overnight incubation, 75 % ethanol was added to the seedlings and left overnight to remove chlorophyll. Arabidopsis seedlings were put on white background and their images were taken to compare T-DNA delivery efficiencies between the auxotrophic and their corresponding WT strains.

### Thymidine-dependent growth of auxotrophs

Thymidine-dependent growth was monitored in a liquid YEP medium. Seed cultures of thymidine auxotrophs (EHA101Thy-, EHA105thy-and EHA105DThy-) and their parental strains were grown in 10 ml of YEP medium in a 50 ml falcon tube in a shaking incubator for 15 hrs at 28°C with 200 rpm. A batch culture was prepared by transferring a calculated amount of overnight culture to a 50 ml of YEP medium in a 250 mL flask to a cell density of 0.02 OD_550_. Batch cultures were grown in a shaking incubator (28°C, 200 prm) and 0.5 mL of culture was sampled every 2 hrs for 24 hrs to measure optical density using a spectrophotometer.

The number of viable cells in the batch culture was monitored for the first 8 hrs. One hundred microliter of a culture was sampled every 2 hrs and serially diluted. One hundred microliter of the diluted cultures (x10^5^ and x10^6^) were spread on solid YEP agar plates supplemented with appropriate antibiotics and 50 mg/L thymidine. Plates were incubated at 28 °C for 48 hrs and the number of colony forming units (CFU/ml) at each time point was determined.

## Results and Discussion

### Generation of thymidine auxotrophs

The overall procedure to generate *thyA* knockout mutants was illustrated in Figure 1. In the first screening of EHA105Thy-mutant, we supplemented the YEP medium with 5 % sucrose and 50 mg/L thymidine. A total of 120 sucrose-tolerant colonies were screened and only one colony exhibited thymidine-dependent growth without the remaining vector backbone. PCR screening amplified an expected 274 bp fragment from the EHA105Thy-colony (1003 bp from the EHA105) suggesting that it was likely a *thyA* knockout mutant (Figure 2A). Similar screening of EHA101Thy-colonies also showed that one *thyA* knockout mutant was obtained (Figure 2B). The low knockout/WT ratio (1/120) was likely attributed to the lethality of the *thyA* mutation within *Agrobacterium* cells. Therefore, we increased thymidine concentration from 50 to 150 mg/L for EHA105DThy-screening, and obtained 6 knockout candidates from the 120 colonies (6/120), suggesting that increased thymidine concentration in the medium can enhance the survival of *thyA* knockout mutant cells after the second recombination. PCR screening confirmed that all six colonies carried a *thyA* deletion (Figure 2C).

**Figure 1.**
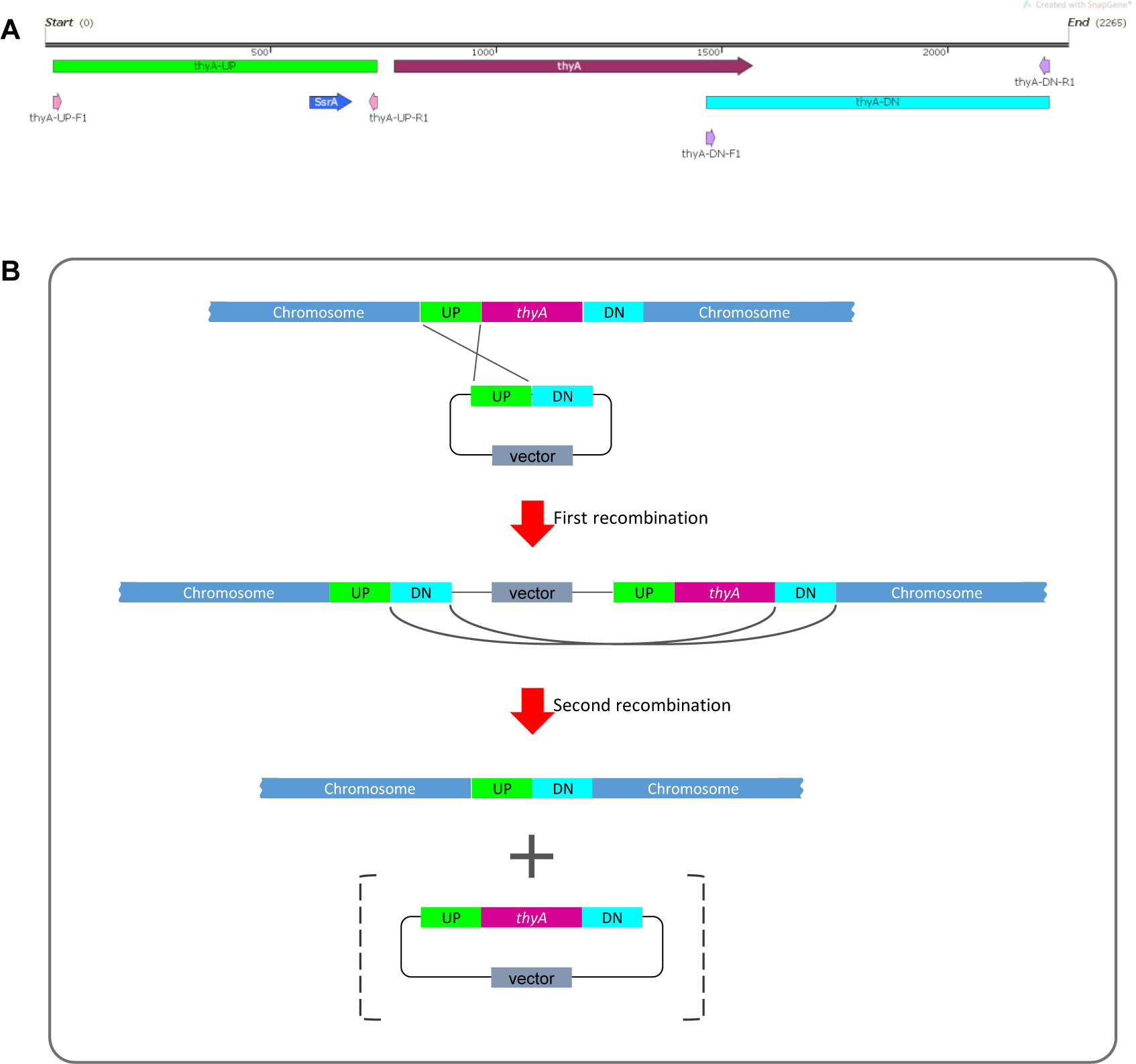
A graphic illustration of homologous recombination mediated *thyA* knockout in *Agrobacterium tumefaciens*. (A) Map of the *thyA* region in the circular chromosome of *A. tumefaciens* C58. The upstream (UP) and downstream (DN) flanking sequences of *thyA* and the primers used for PCR are indicated. A transfer-messenger RNA (tmRNA), SsrA, was part of the UP flanking sequence. (B) Single homologous recombination between the upstream (or downstream) flanking sequences leads to integration of the knockout construct into the *Agrobacterium* chromosome. Antibiotics resistance gene encoded on the vector backbone confers resistance during bacterial selection. Antibiotic resistant *Agrobacterium* cells are sensitive to sucrose due to the *sacB* gene encoded on the vector backbone, whose product converts sucrose into levan, a toxic molecule (Gay et al., 1983; Steinmetz et al., 1983). During the negative selection on the antibiotics-free medium containing 5% sucrose and 150 mg/L of thymidine, second homologous recombination between the downstream flanking sequences leads to deletion of the vector backbone and *thyA* gene. Sucrose-tolerant *Agrobacterium* cells are tested for antibiotics sensitivity and thymidine dependence growth. PCR screening is used to identify *thyA* knockout mutants, which are sensitive to antibiotics and thymidine dependent.

**Figure 2.**
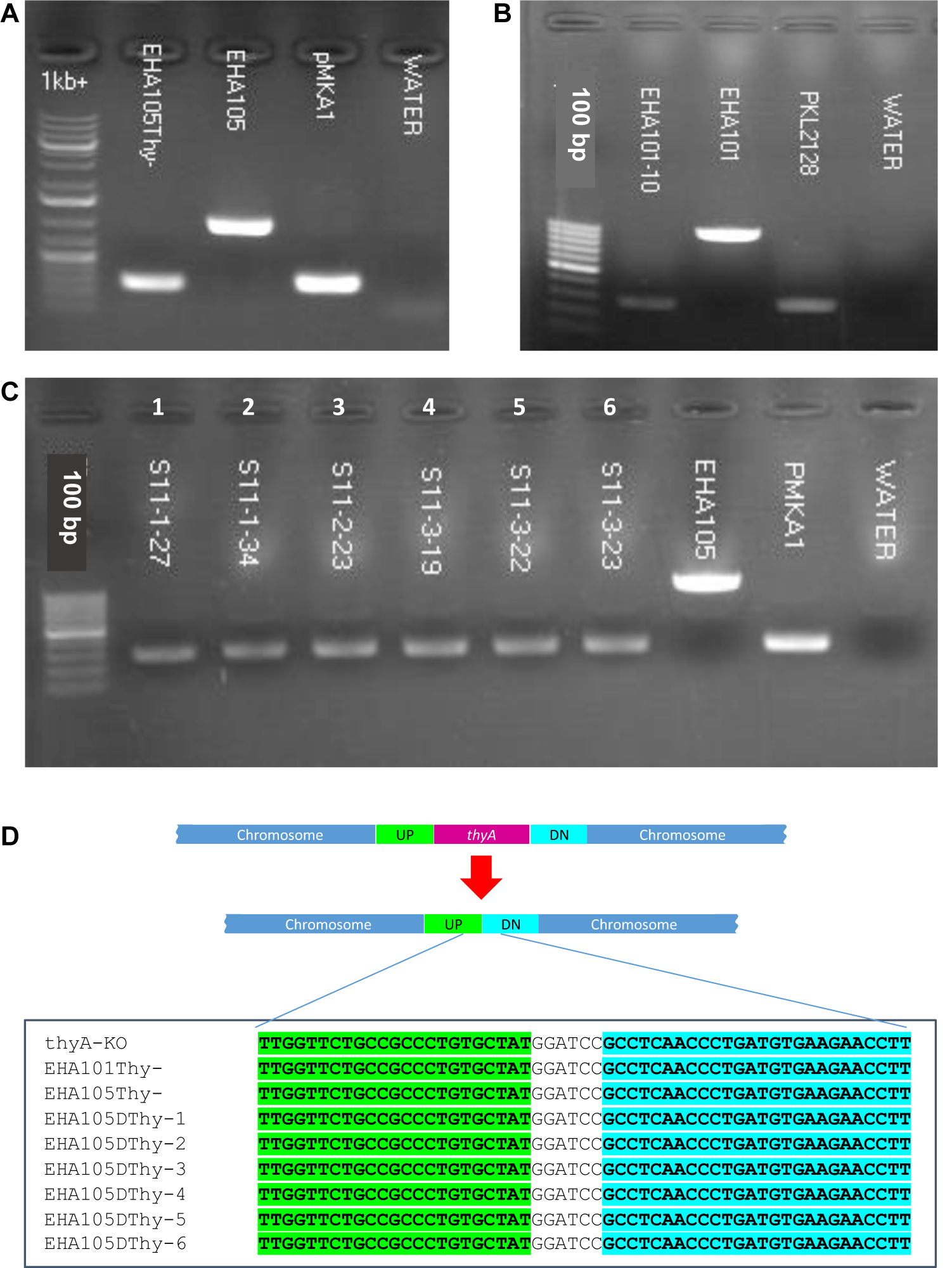
Screening of *thyA* knockout mutants by PCR and Sanger sequencing. (A) EHA105Thy-, (B) EHA101, (C) EHA105DThy-, and (D) Multiple sequence alignment of the Sanger sequencing results of the *thyA* deletion junction. EHA101 or EHA105 were used as WT control (1003 bp), while pMKA1 was used as a knockout control (274 bp). The junction of the UP and DN flanking sequences contains the *Bam*HI recognition site (GGATCC) introduced during the knockout vector construction.

The junction sequences of the *thyA* knockout mutants were subjected to Sanger sequencing to verify the precise sequence deletion by homologous recombination. Sequencing results showed that all knockout mutants carried intended *thyA* deletion mutation (Figure 2D). Interestingly, however, three of the six EHA105DThy-knockout mutants (Figure 2D, EHA105DThy-4, -5, and -6) carried additional 10 bp deletion within the coding sequence of an upstream gene, *Atu2049*, which encodes a transfer-messenger RNA (tmRNA) SsrA (Figure S1C); therefore, we removed these mutant strains from further analyses. It is not clear whether these three mutants were clonal or not; but the presence of additional mutation in the *thyA* flanking region indicates that even homologous recombination mediated gene knockout approach can result in unintended mutations, therefore close examination of the junction regions are necessary to avoid unnecessary complication of the downstream analyses.

### Examination of T-DNA transfer capability

We next tested if the thymidine auxotrophic strains can efficiently deliver T-DNAs into plant cells using the AGROBEST assay (Wu et al., 2014). As shown in Figure 3, GUS staining results demonstrated that all three auxotrophic strains retain the T-DNA delivery capability. Both EHA101Thy-(Figure 3A) and EHA105Thy-(Figure 3C) strains showed similar level of GUS expression in the Arabidopsis seedlings compared to their corresponding prototrophs (Figure 3B and 3D, respectively) and the reference strain LBA4404Thy-(Figure 3 G), whereas EHA105DThy-strain (Figure 3E) showed relatively weaker GUS expression compared to EHA105D (Figure 3F). Further study is needed to determine if the seemingly weaker GUS expression is an indication of diminished T-DNA transfer capability of the EHA105DThy-strain and if it carries additional mutations other than *thyA* deletion in the genome. In sum, EHA101Thy-, EHA105Thy- and EHA105DThy-strains can deliver T-DNA into Arabidopsis cells and they are ready to be used for transient and stable plant transformation applications.

**Figure 3.**
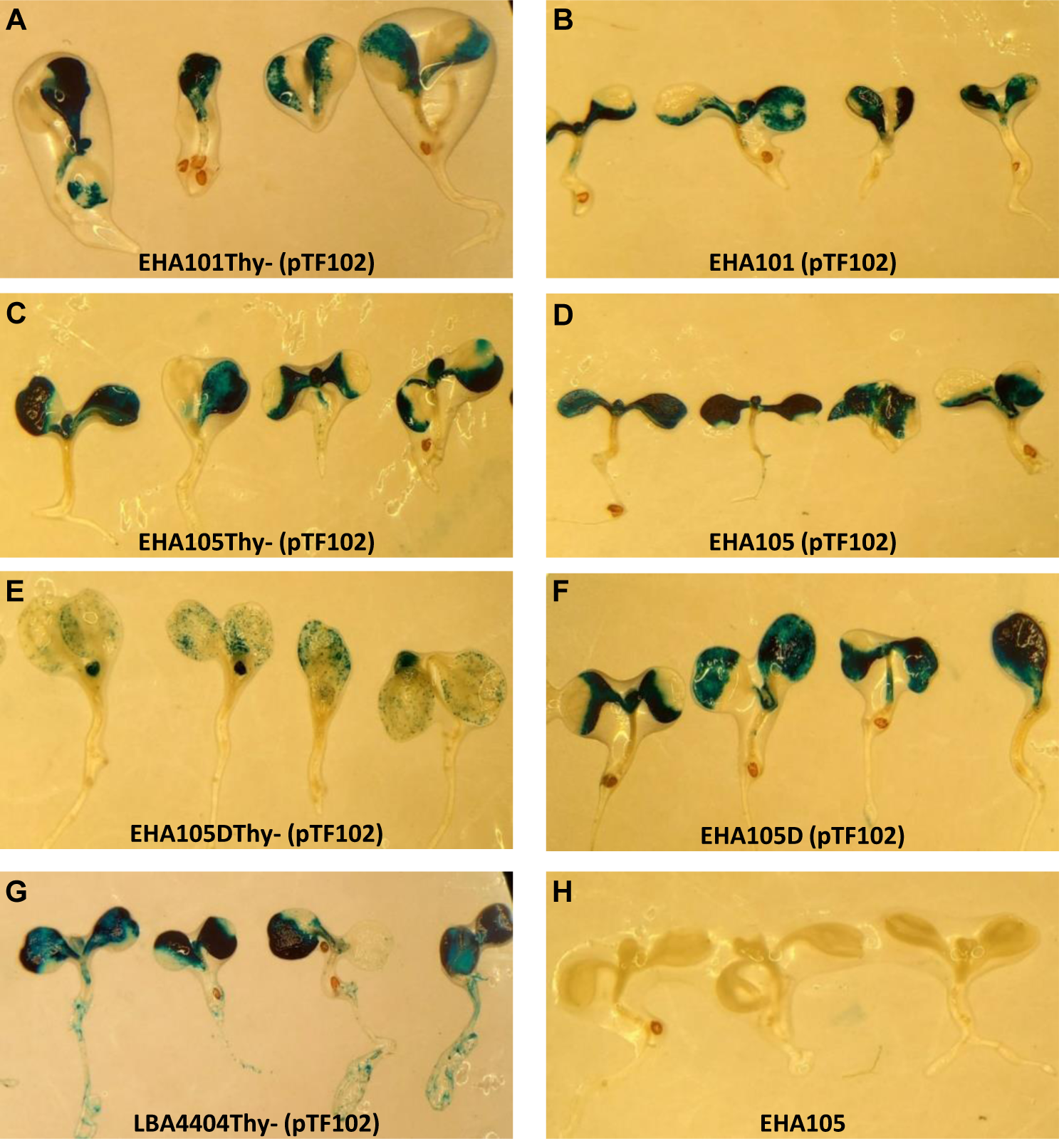
Transient GUS expression assay (AGROBEST) using Arabidopsis *efr-1* seedlings by various *Agrobacterium* strains. Seven-day-old Arabidopsis *efr-1* seedlings infected with different prototrophic and thymidine auxotrophic *Agrobacterium* stains carrying the binary vector pTF102 were compared by GUS staining. EHA105 strain without pTF102 (H) served as a negative control.

### Thymidine-dependent growth of auxotrophic *Agrobacterium* strains

As all three auxotrophic strains were selected based on their lack of growth on YEP medium without thymidine supplement, they did show thymidine-dependent growth in liquid medium. As mentioned above, increasing thymidine concentration from 50 mg/L to 150 mg/L was helpful to recover more *thyA* knockout mutants (1/120 vs. 6/120). In addition to this, we monitored the growth of each strain in liquid YEP medium supplemented with three different concentrations of thymidine as well as appropriate antibiotics. As expected, all tested auxotrophic strains showed increased growth rate and maintained higher cell density when supplemented with higher concentrations of thymidine (Figure 4). Interestingly, the prototrophic strains grew faster than the auxotrophic strains even in the presence of 150 mg/L thymidine suggesting that thymidine uptake might be a limiting factor for the auxotrophic strains. Compared to other strains, EHA101 and EHA101Thy-grew slightly slower, presumably due to the presence of kanamycin in addition to spectinomycin in the medium for other strains. Overall, the average cell density of the auxotrophic strains across the three different thymidine concentrations (Figure 4E) summarized the growth pattern: EHA101Thy-grew slightly slower than EHA105Thy- and EHA105DThy-, which showed nearly identical growth rate over the time course. Lastly, we monitored the number of viable cells from the liquid cultures during the first 8 hrs of growth (Figure 5). There was no noticeable difference in the relationship between the optical cell density and the number of viable cells among the auxotrophic and prototrophic strains, further suggesting that these thymidine auxotrophic strains can be properly grown and maintained by supplementing 50-150 mg/L thymidine.

**Figure 4.**
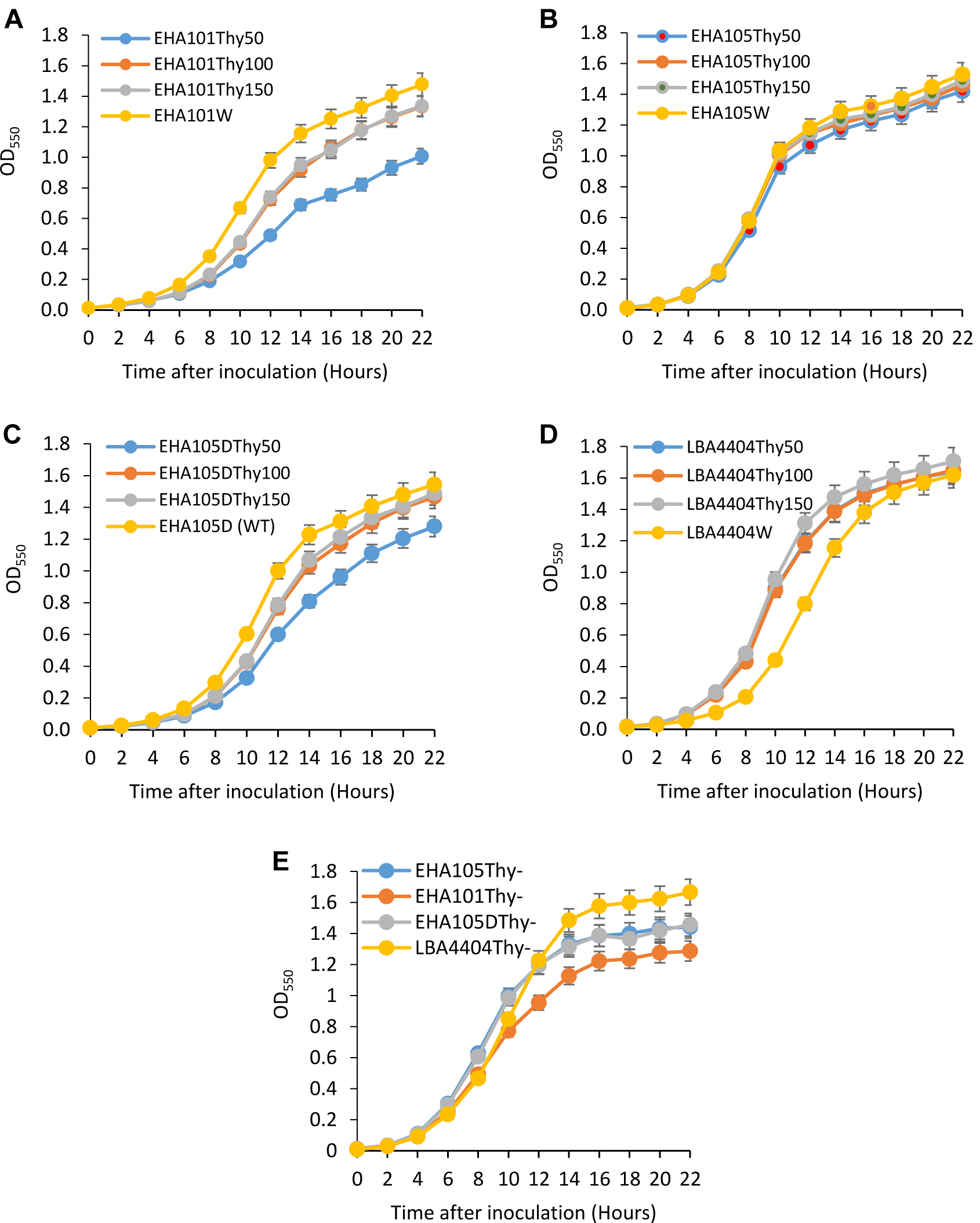
Thymidine-dependent growth of auxotrophic *Agrobacterium* strains. Optical density of *Agrobacterium* cells grown with varying amount of thymidine were monitored for 22 hrs at 550 nm using a spectrophotometer: growth curves of WT (yellow curve) and thymidine auxotrophs (blue, ash, and orange curves) with thymidine concentrations of 50 mg/L (Thy50), 100 mg/L (Thy100), and 150 mg/L (Thy150). (A) EHA101 and EHA101Thy-, (B) EHA105 and EHA105Thy-, (C) EHA105D and EHA105DThy-, (D) LBA4404 and LBA4404Thy-, (E) Average cell density of *Agrobacterium* thymidine auxotrophs with 50 mg/L of thymidine: EHA101Thy-(orange curve), EHA105Thy-(blue curve), EHA105DThy-(ash curve), and LBA4404Thy-(yellow curve). Data represent mean (± standard deviation) of three replicates.

**Figure 5.**
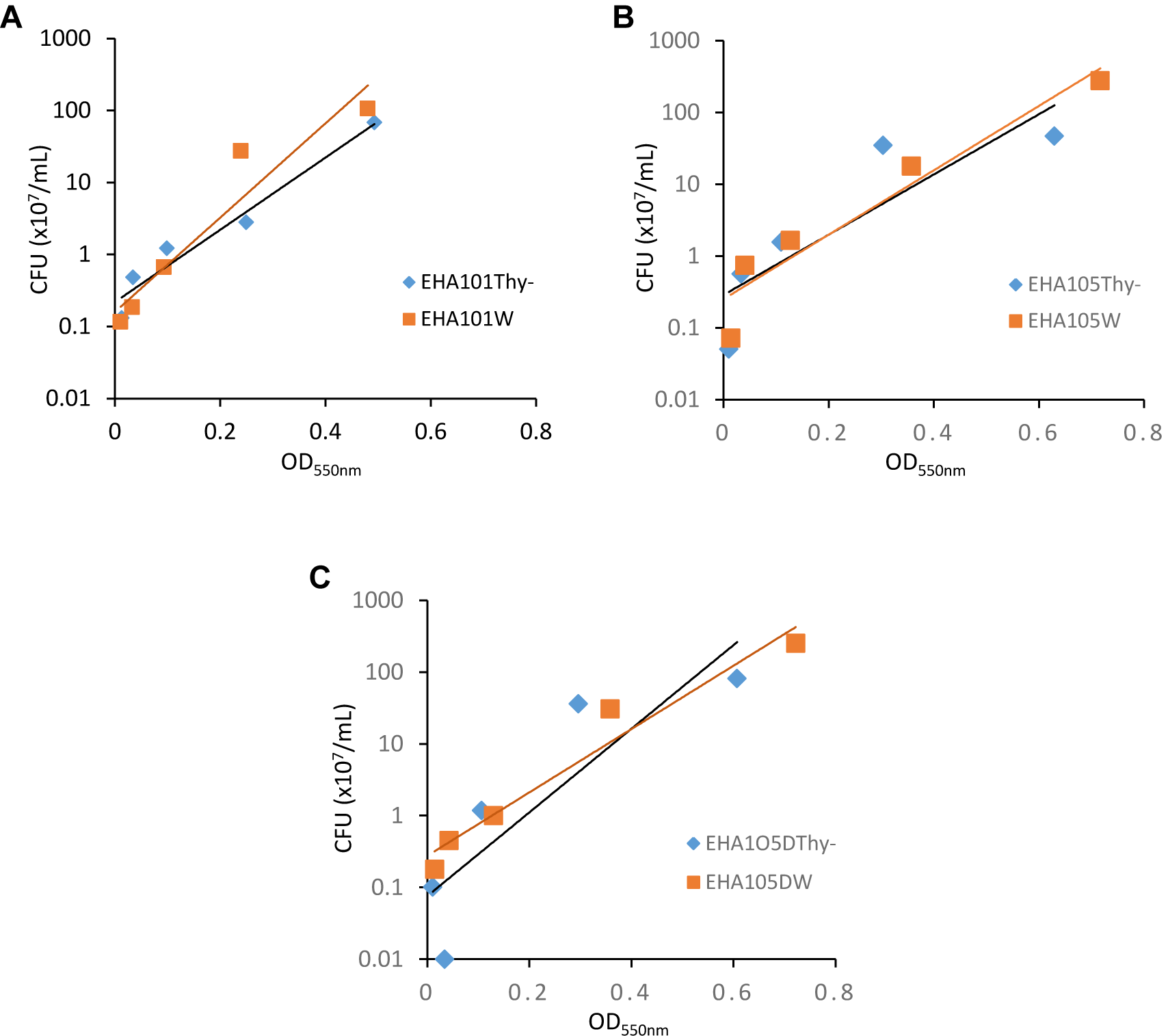
Correlation between the numbers of viable cells (CFU/mL) and optical cell density (OD_550_). The number of viable cells and optical cell density were measured every two hours for the first 8 hours of growth and the trendlines depict a high level of correlation between them. (A) EHA101 and EHA101Thy-, (B) EHA105 and EHA105Thy-, and (C) EHA105D and EHA105DThy-.

## Conclusion

We successfully generated thymidine auxotrophic strains from the commonly used *Agrobacterium* strains EHA101 and EHA105 for plant transformation. Because *thyA* gene was completely deleted via homologous recombination, these strains are stably thymidine-dependent hence can be effectively eliminated from the explants after co-cultivation by omitting thymidine from the medium. Importantly, these auxotrophic strains can deliver T-DNAs into plant cells as effectively as their prototrophs, thus they are ready for plant transformation applications. These auxotrophic strains are available upon request.

## Acknowledgements

We thank Corteva Agriscience for providing the LBA4404Thy-strain. This project was partially supported by National Science Foundation Plant Genome Research Program Grant 1725122 and 1917138 to K.W., by the Iowa State University Interdepartmental Plant Biology Major fellowship to EA and MA, by the USDA NIFA Hatch project #IOW04341 and by State of Iowa funds and by the Crop Bioengineering Center of Iowa State University.

## Supplementary Information

### Supplementary Table

**Table S1. Media composition**

### Supplementary Figure legend

**Figure S1. Maps of the *thyA* knockout constructs and sequence alignment of *thyA* junction sequences**. (A) pMKA1 has a kanamycin resistance gene and was used for EHA105 and EHA105D. (B) pKL2128 has a spectinomycin resistance gene and was used for EHA101. (C) Multiple sequence alignment of the Sanger sequencing results of the *thyA* UP flanking sequence near the junction. Three mutants (EHA105DThy-4, 5, 6) had a 10 bp deletion within the coding sequence of the *Atu2049*, which encodes a transfer-messenger RNA (tmRNA), SsrA.

## References

Anand A., Bass S. H., Wu E., Wang N., McBride K. E., Annaluru N., ….., and Jones T. J. (2018). An improved ternary vector system for Agrobacterium-mediated rapid maize transformation. Plant Mol Biol 97: 187–200.

Britton M. T., Escobar M. A., and Dandekar A. M. (2008). The oncogenes of Agrobacterium tumefaciens and Agrobacterium rhizogenes. In: Agrobacterium (Tzfira T. and Citovsky V., eds.). Springer, New York, NY, pp. 524–563.

Cervera M. (2005). Histochemical and fluorometric assays for uidA (GUS) gene detection. In: Transgenic Plants: Methods and Protocols (Peña L., ed). Humana Press Inc, Totowa, NJ, pp 203–213. https://doi.org/10.1385/1-59259-827-7:203

Frame B. R., Shou H., Chikwamba R. K., Zhang Z., Xiang C., Fonger T. M., ……, and Wang K. (2002). Agrobacterium tumefaciens-mediated transformation of maize embryos using a standard binary vector system. Plant Physiol 129: 13–22.

Gay P., Le Coq D., Steinmetz M., Ferrari E., and Hoch J. A. (1983). Cloning structural gene sacB, which codes for exoenzyme levansucrase of Bacillus subtilis: expression of the gene in Escherichia coli. J Bacteriol 153: 1424–1431.

Gelvin S. B. (2003). Agrobacterium-mediated plant transformation: The biology behind the “gene-jockeying” tool. Microbiol Mol Biol Rev 67: 16–37.

Gelvin S. B. (2006). Agrobacterium virulence gene induction. In: Agrobacterium Protocols, 2^nd^ edition (Wang K, ed). Humana Press Inc, Totowa, NJ, pp 77–84.

Gelvin S. B. (2017). Integration of Agrobacterium T-DNA into the plant genome. Annu Rev Genet 51: 195–217.

Hood E. E., Gelvin S. B., Melchers L. S., and Hoekema A. (1993). New Agrobacterium helper plasmids for gene transfer to plants. Transgenic Res 2: 208–218.

Hood E. E., Helmer G. L., Fraley R. T., and Chilton M. D. (1986). The hypervirulence of Agrobacterium tumefaciens A281 is encoded in a region of pTiBo542 outside of T-DNA. J Bacteriol 168: 1291–1301.

Kim Y-G., Sharmin S. A., Alam I., Kim K-H., Kwon S-Y., Sohn J-H., ……, and Lee B-H. (2013). Agrobacterium-mediated transformation of reed (Phragmites communis Trinius) using mature seed-derived calli. GCB Bioenergy 5: 73–80.

Komari T., Halperin W., and Nester E. W. (1986). Physical and functional map of supervirulent Agrobacterium tumefaciens tumor-inducing plasmid pTiBo542. J Bacteriol 166: 88–94.

Komari T., Takakura Y., Ueki J., Kato N., Ishida Y., and Hiei Y. (2006). Binary vectors and super-binary vectors. In: Agrobacterium Protocols, 2^nd^ edition (Wang K, ed). Humana Press Inc, Totowa, NJ, pp 15–41.

Koncz C. and Schell J. (1986). The promoter of TL-DNA gene 5 controls the tissue-specific expression of chimaeric genes carried by a novel type of Agrobacterium binary vector. Mol Gen Genet 204: 383–396.

Lazo G. R., Stein P. A., and Ludwig R. A. (1991). A DNA transformation-competent Arabidopsis genomic library in Agrobacterium. Biotechnol 9: 963–967.

Lee L. Y. and Gelvin S. B. (2008). T-DNA binary vectors and systems. Plant Physiol 146: 325–332.

Lee K., Huang X., Yang C., Lee D., Ho V., Nobuta K., ……, and Wang K. (2013). A genome-wide survey of highly expressed non-coding RNAs and biological validation of selected candidates in Agrobacterium tumefaciens. PLoS One 8: e70720.

Lowder L. G., Paul J. W., Baltes N. J., Voytas D. F., Zhang Y., Zhang D., ……, and Qi Y. (2015). A CRISPR/Cas9 toolbox for multiplexed plant genome editing and transcriptional regulation. Plant Physiol 169: 971–985.

Mattanovich D., Rüker F., Machado A. C., Laimer M., Regner F., Steinkellner H., ……, and Katinger H. (1989). Efficient transformation of Agrobacterium spp. by electroporation. Nucleic Acids Res. 17: 6747.

Matthysse A. G., Yarnall H., Boles S. B., and McMahan S. (2000). A region of the Agrobacterium tumefaciens chromosome containing genes required for virulence and attachment to host cells. Biochim Biophys Acta 1490: 208–212.

Nauerby B., Billing K., and Wyndaele R. (1997). Influence of the antibiotic timentin on plant regeneration compared to carbenicillin and cefotaxime in concentration suitable for elimination of Agrobacterium tumefaciens. Plant Science 123: 169–177.

Ooms G., Hooykaas P. J. J., Moolenaar G., and Schilperoort R. A. (1981). Crown gall plant tumors of abnormal morphology, induced by Agrobacterium tumefaciens carrying mutated octopine Ti plasmids; analysis of T-DNA functions. Gene 14: 33–50.

Pollock K., Barfield D. G. and Shields R. (1983). The toxicity of antibiotics to plant cell cultures. Plant Cell Rep 2: 36–39.

Ranch J. P., Liebergesell M., Garnaat C. W. and Huffman G. A. (2010). Auxotrophic Agrobacterium for plant transformation and methods thereof. WO application WO 2010078445A1.

Schäfer A., Tauch A., Jäger W., Kalinowski J., Thierbach G., and Puhler A. (1994). Small mobilizable multi-purpose cloning vectors derived from the Escherichia coli plasmids pK18 and pK19: selection of defined deletions in the chromosome of Corynebacterium glutamicum. Gene 145: 69–73.

Sciaky, D., Montoya, A. L. and Chilton, M. D. (1978). Fingerprints of Agrobacterium Ti plasmids. Plasmid 1: 238–253.

Steinmetz M., Le Coq D., Djemia H. B., and Gay P. (1983). Genetic analysis of sacB, the structural gene of a secreted enzyme, levansucrase of Bacillus subtilis Marburg. Mol Gen Genet 191: 138–144.

Tempé J. and Petit A. (1982). Opine utilization by Agrobacterium In: Molecular Biology of Plant Tumors (Kahl G. and Schell J, eds). Academic Press, New York, NY, pp. 451–459.

Tzfira T. and Citovsky V. (2006). Agrobacterium-mediated genetic transformation of plants: Biology and biotechnology. Curr Opin Biotech 17: 147–154.

Untergasser A., Cutcutache I., Koressaar T., Ye J., Faircloth B. C., ……, and Rozen S. G. (2012). Primer3-new capabilities and interfaces. Nucleic Acids Res 40: 3115.

Wu H., Liu K., Wang Y., Wu J. F., Chiu W. L., Chen C. Y., ……, and Lai E. M. (2014). AGROBEST: an efficient Agrobacterium-mediated transient expression method for versatile gene function analyses in Arabidopsis seedlings. Plant Methods 10: 19.

