## Supplementary material for "Generation of thymidine auxotrophic *Agrobacterium tumefaciens* strains for plant transformation": Table S1

### Table S1A. YEP medium.

| Ingredient | Quantity per liter (L) |
|---|---|
| Yeast Extract | 10 g |
| Bacto Peptone | 10 g |
| Sodium chloride (NaCl) | 5 g |
| Agar (solid medium only) | 15 g |

### Table S1B. *Agrobacterium* induction medium (Gelvin 2006).

| 20 x AB salts | Quantity per liter (L) |
|---|---|
| Ammonium chloride ($NH_4Cl$) | 20 g |
| Magnesium sulfate ($MgSO_4 \cdot 7H_2O$) | 6 g |
| Potassium chloride (KCl) | 3 g |
| Calcium chloride ($CaCl_2$) | 0.2 g |
| Ferrous sulfate ($FeSO_4 \cdot 7H_2O$) | 50 mg |
| **Induction medium** | **Final conc.** |
| AB salts | 1x |
| Trisodium phosphate ($Na_3PO_4$) | 2 mM |
| MES, pH 5.6 | 50 mM |
| Glucose | 0.5% |
| Acetosyringone | 100 µM |

### Table S1C. MS medium composition (Murashige and Skoog, 1962).

| Major salts (macronutrients) | Quantity per liter (L) |
|---|---|
| Ammonium nitrate ($NH_4NO_3$) | 1,650 mg |
| Calcium chloride ($CaCl_2 \bullet 2H_2O$) | 440 mg |
| Magnesium sulfate ($MgSO_4 \bullet 7H_2O$) | 370 mg |
| Monopotassium phosphate ($KH_2PO_4$) | 170 mg |
| Potassium nitrate ($KNO_3$) | 1900 mg |
| **Minor salts (micronutrients)** | **Quantity per liter (L)** |
| Boric acid ($H_3BO_3$) | 6.2 mg |
| Cobalt chloride ($CoCl_2 \bullet 6H_2O$) | 0.025 mg |
| Ferrous sulfate ($FeSO_4 \bullet 7H_2O$) | 27.8 mg |
| Manganese(II) sulfate ($MnSO_4 \bullet 4H_2O$) | 22.3 mg |
| Potassium iodide (KI) | 0.83 mg |
| Sodium molybdate ($Na_2MoO_4 \bullet 2H_2O$) | 0.25 mg |
| Zinc sulfate ($ZnSO_4 \bullet 7H_2O$), 8.6 mg/L | 8.6 mg |
| Ethylenediaminetetraacetic acid ferric sodium (NaFe-EDTA) containing **5.57 g** $FeSO_4 \bullet 7H_2O$ and **7.45 g** $Na_2$-EDTA per litre of water | 5 ml |
| Copper sulfate ($CuSO_4 \bullet 5H_2O$) | 0.025 mg |
| **Vitamins and organic compounds** | **Quantity per liter (L)** |
| Myo-Inositol | 100 mg |
| Nicotinic Acid | 0.5 mg |
| Pyridoxine · HCl | 0.5 mg |
| Thiamine · HCl | 1.0 mg |
| Glycine | 2 mg |

### Supplemental References


Gelvin S. B. (2006) *Agrobacterium* virulence gene induction. In: Agrobacterium Protocols, 2nd edition (Wang K, ed). Humana Press Inc, New Jersey, USA. pp 77–84.

Murashige T. and Skoog F. (1962). A Revised Medium for Rapid Growth and Bio Assays with Tobacco Tissue Cultures. *Physiologia Plantarum*. 15 (3): 473–497.
