## Supplementary material for "Generation of thymidine auxotrophic *Agrobacterium tumefaciens* strains for plant transformation": Figure S1

**A**

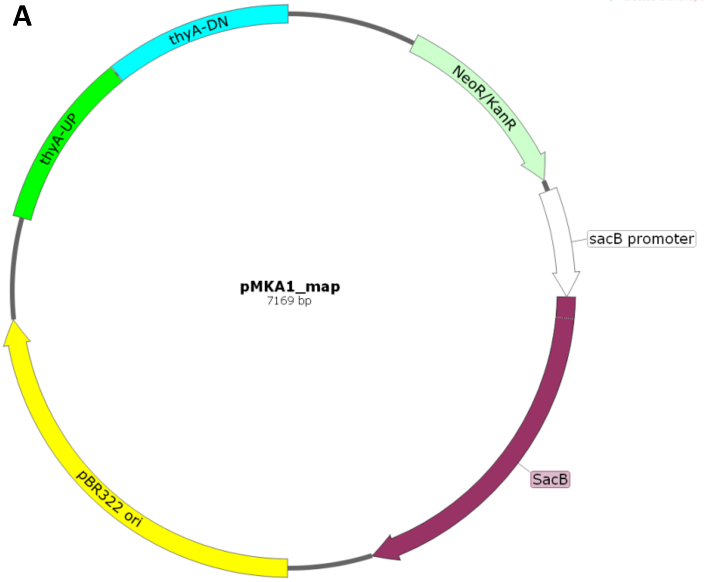


**B**

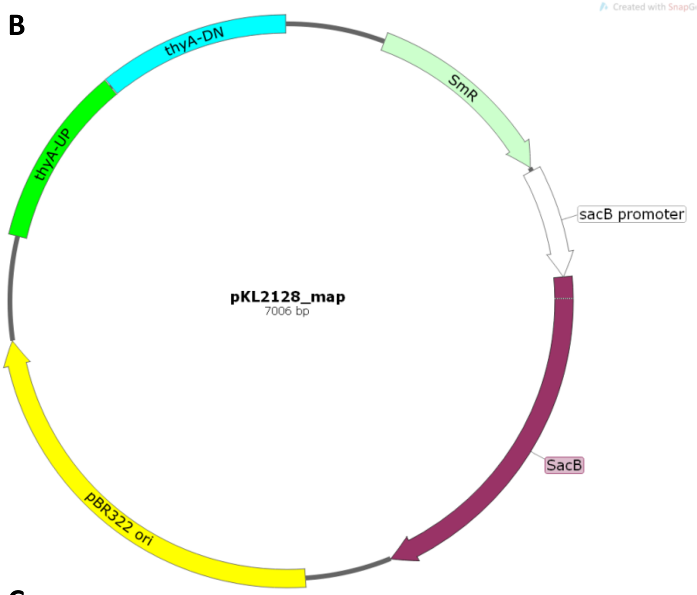


**C**

```
                                                                    Atu2049
                                                                      Stop
thyA-KO          CTTCTATTATCTGGTGGCCAAGGAACTGGGCAAGCTTCAGTCGCTTTGATCTGAG
EHA101Thy-       CTTCTATTATCTGGTGGCCAAGGAACTGGGCAAGCTTCAGTCGCTTTGATCTGAG
EHA105Thy-       CTTCTATTATCTGGTGGCCAAGGAACTGGGCAAGCTTCAGTCGCTTTGATCTGAG
EHA105DThy-1     CTTCTATTATCTGGTGGCCAAGGAACTGGGCAAGCTTCAGTCGCTTTGATCTGAG
EHA105DThy-2     CTTCTATTATCTGGTGGCCAAGGAACTGGGCAAGCTTCAGTCGCTTTGATCTGAG
EHA105DThy-3     CTTCTATTATCTGGTGGCCAAGGAACTGGGCAAGCTTCAGTCGCTTTGATCTGAG
EHA105DThy-4     CTTCTATTATCTGGTGGCCA----------CAAGCTTCAGTCGCTTTGATCTGAG
EHA105DThy-5     CTTCTATTATCTGGTGGCCA----------CAAGCTTCAGTCGCTTTGATCTGAG
EHA105DThy-6     CTTCTATTATCTGGTGGCCA----------CAAGCTTCAGTCGCTTTGATCTGAG
                 ********************          *************************
```

**Figure S1.** Maps of the *thyA* knockout constructs and sequence alignment of *thyA* junction sequences. (A) pMKA1 has a kanamycin resistance gene and was used for EHA105 and EHA105D. (B) pKL2128 has a spectinomycin resistance gene and was used for EHA101. (C) Multiple sequence alignment of the Sanger sequencing results of the *thyA* UP flanking sequence near the junction. Three mutants (EHA105DThy-4, 5, 6) had a 10 bp deletion within the coding sequence of the *Atu2049*, which encodes a transfer-messenger RNA (tmRNA), SsrA.
